# Searching through functional space reveals distributed visual, auditory, and semantic coding in the human brain

**DOI:** 10.1101/2020.04.20.052175

**Authors:** Sreejan Kumar, Cameron T. Ellis, Thomas O’Connell, Marvin M Chun, Nicholas B. Turk-Browne

## Abstract

The extent to which brain functions are localized or distributed is a foundational question in neuroscience. In the human brain, common fMRI methods such as cluster correction, atlas parcellation, and anatomical searchlight are biased by design toward finding localized representations. Here we introduce the functional searchlight approach as an alternative to anatomical searchlight analysis, the most commonly used exploratory multivariate fMRI technique. Functional searchlight removes any anatomical bias by grouping voxels based only on functional similarity and ignoring anatomical proximity. We report evidence that visual and auditory features from deep neural networks and semantic features from a natural language processing model are more widely distributed across the brain than previously acknowledged. This approach provides a new way to evaluate and constrain computational models with brain activity and pushes our understanding of human brain function further along the spectrum from strict modularity toward distributed representation.

## Introduction

One of the most important debates throughout the history of neuroscience has been whether each mental function is localized to a dedicated brain region or distributed across regions[1]. Early studies of patients with specific brain damage and accompanying behavioral deficits suggested that some functions, such as language and executive control, can be localized. This localist perspective provided the foundation for initial studies of the healthy brain with functional magnetic resonance imaging (fMRI), which identified regions of interest with circumscribed functions[2]. Subsequent studies, however, showed support for a distributional perspective by suggesting that some functions instead arise out of the joint action of multiple regions[3]. Such claims were supported by the emergence of multivariate methods that decode the function of patterns of fMRI activity across populations of voxels[4].

Despite the promise of multivariate methods, the predominant exploratory approach for finding distributed representations with them remains inherently localist. Specifically, patterns of activity are extracted from small, contiguous anatomical volumes by moving a cube or sphere of voxels, known as a “searchlight”, throughout the brain[5]. These patterns of activity are passed on for subsequent multivariate analysis, such as decoding the category of a stimulus. Although a valuable tool for mapping the informational contents of individual regions, information spread across disparate regions is never included within the same searchlight and thus neglected. One potential solution is to extract whole-brain patterns of activity[6]. However, even if the information is distributed throughout the brain, only a fraction of voxels would be expected to contribute, hence these multivariate models would be hard to fit and suffer from the curse of dimensionality. Even if successful, it is hard to interrogate the model to determine how the information is represented, because it is possible to observe different voxel weights for identical activity profiles[7].

We hypothesized that for many cognitive functions, information is widely distributed throughout the brain and that evidence for localized representation may partially be attributed to anatomical constraints of current methods. To test this hypothesis, we developed a new searchlight approach that is not bound by anatomy. This *functional* searchlight retains the exhaustive search of traditional anatomical searchlight while eliminating the localist assumption that only neighboring voxels contain useful information. Specifically, we re-map voxels from their original 3-D anatomical space with coordinates in *x* (left-right), *y* (anterior-posterior), and *z* (inferior-superior) dimensions into a new functional space with dimensions for orthogonal latent variables that capture reliable sources of variation in brain activity irrespective of anatomy (Figure 1).

**Figure 1.**
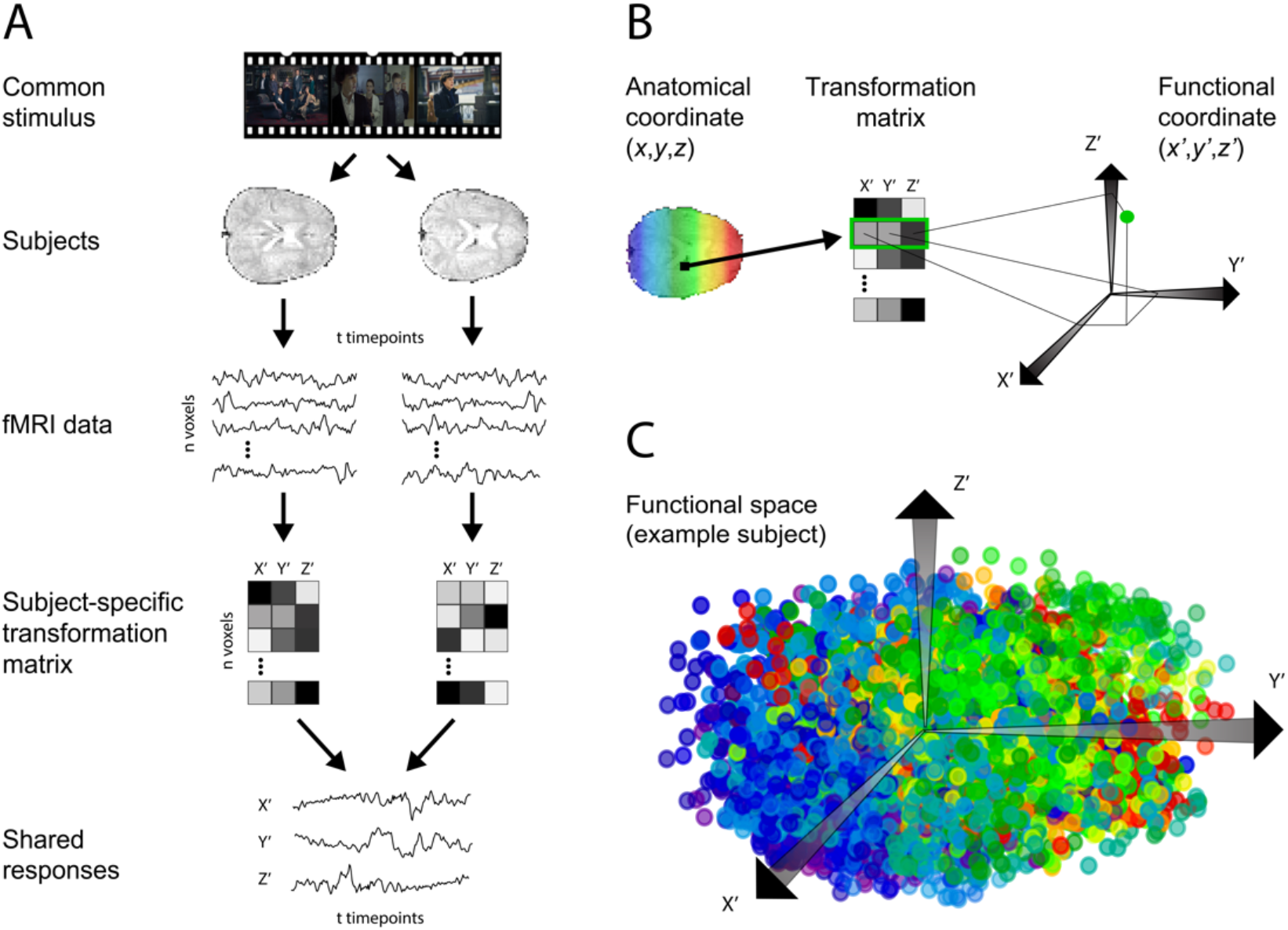
Transformation from anatomical space to functional space. (A) We use shared response modeling (SRM) to transform whole-brain data into a *k*-dimensional shared feature space. (B) Each voxel is transformed into functional space by using its loadings on the dimensions of the shared space as its coordinates in functional space. (C) Example functional space for one subject. Color indicates position on the anterior-posterior *y* axis of the input anatomical space, and as can be seen voxels get reorganized in functional space. Voxels that are functionally similar but anatomically disparate can be grouped together (e.g., blue-purple and red in top left). Note that this functional space is three-dimensional for visualization purposes, but the functional space used in our analyses had 200 dimensions.

The functional space was learned through shared response modeling (SRM)[8], which finds a *k*-dimensional representation capturing the information shared across participants viewing the same stimulus. In functional space, two voxels are arranged close together if their projection weights into the shared space are similar, that is, if they load similarly on the latent variables. Because proximity in the functional space is governed entirely by this correspondence and not by anatomical distance, a single searchlight in this space could have access to information that is anatomically distributed throughout the brain. To the extent that activity patterns extracted from these functional searchlights better represent task information than those from anatomical searchlights over contiguous voxels, the information can be said to be distributed.

## Results and Discussion

To fit the SRM and compare the performance of functional vs. anatomical searchlights, we used an fMRI dataset in which participants watched an episode of BBC’s “Sherlock”[9]. SRM requires a hyperparameter for the number of dimensions, which we determined to be *k*=200 by performing time-segment matching (Figure S1). After learning this functional space on half of the movie, we evaluated how well patterns of activity from functional searchlights encoded visual, auditory, and semantic features using model-based analysis on the other half of the movie. For each functional searchlight, the nearest 342 voxels in this high dimensional space were used to define the searchlight, hence using the same number of voxels as was used in the anatomical searchlight.

We first tested whether voxels in functional searchlights, relative to anatomical searchlights, share more information with representations in stimulus-computable models of visual and auditory processing. Deep neural networks (DNNs) offer a way to extract features from an audio-visual stimulus and measure the expression of these features in the brain[10]. We computed DNN activity separately for the video and audio components of the movie. For video, DNN activity was computed for individual frames using AlexNet[11], a DNN model pre-trained to recognize objects from natural images; hidden layer activity was averaged across video frames that fell within the same TR (1.5s). For audio, 1.5s audio segments were fed into a branching music-speech recognition DNN[12] (referred to here as KellNet).

For each layer in both the visual and auditory networks, a representational similarity matrix (RSM) was computed as the correlation matrix of the DNN activity across all time-points. For a given layer of the network, the pattern of activity across units at every time-point was correlated with every other time-point in the movie. Likewise, an fMRI RSM was computed for each searchlight as the correlation matrix of the BOLD activity patterns in the searchlight across all time-points. The upper triangular elements of these model and brain RSMs were then correlated in a second-order analysis to assess the similarity of the information captured in the fMRI searchlight and DNN layer of a given network. To minimize the contribution of the auto-correlation inherent in these data that would inflate the similarity, a buffer of 10 TRs (15s) was excluded off the diagonal and ignored for subsequent analysis. This procedure was completed for all subjects, searchlight locations, and hidden layers in both the visual and auditory DNNs.

Not all searchlights were expected to represent the audio-visual content of movie. Hence, we compared the performance of the top 1% of functional and anatomical searchlights. The functional searchlight showed a significantly higher RSA for all of the AlexNet layers except the first and for all of the KellNet layers, compared to the anatomical searchlight (see 95% confidence intervals in Figure S2). Improvements for two example layers (conv2 in AlexNet and fc7W in KellNet) are shown in Figure 2A (left). Voxels that consistently contributed to the top 1% of functional searchlights were more distributed than those that consistently contributed to the top 1% of anatomical searchlights (Figure 2B left). By removing the assumption that information is anatomically local, we found neural representations that are more consistently correlated with model representations.

**Figure 2.**
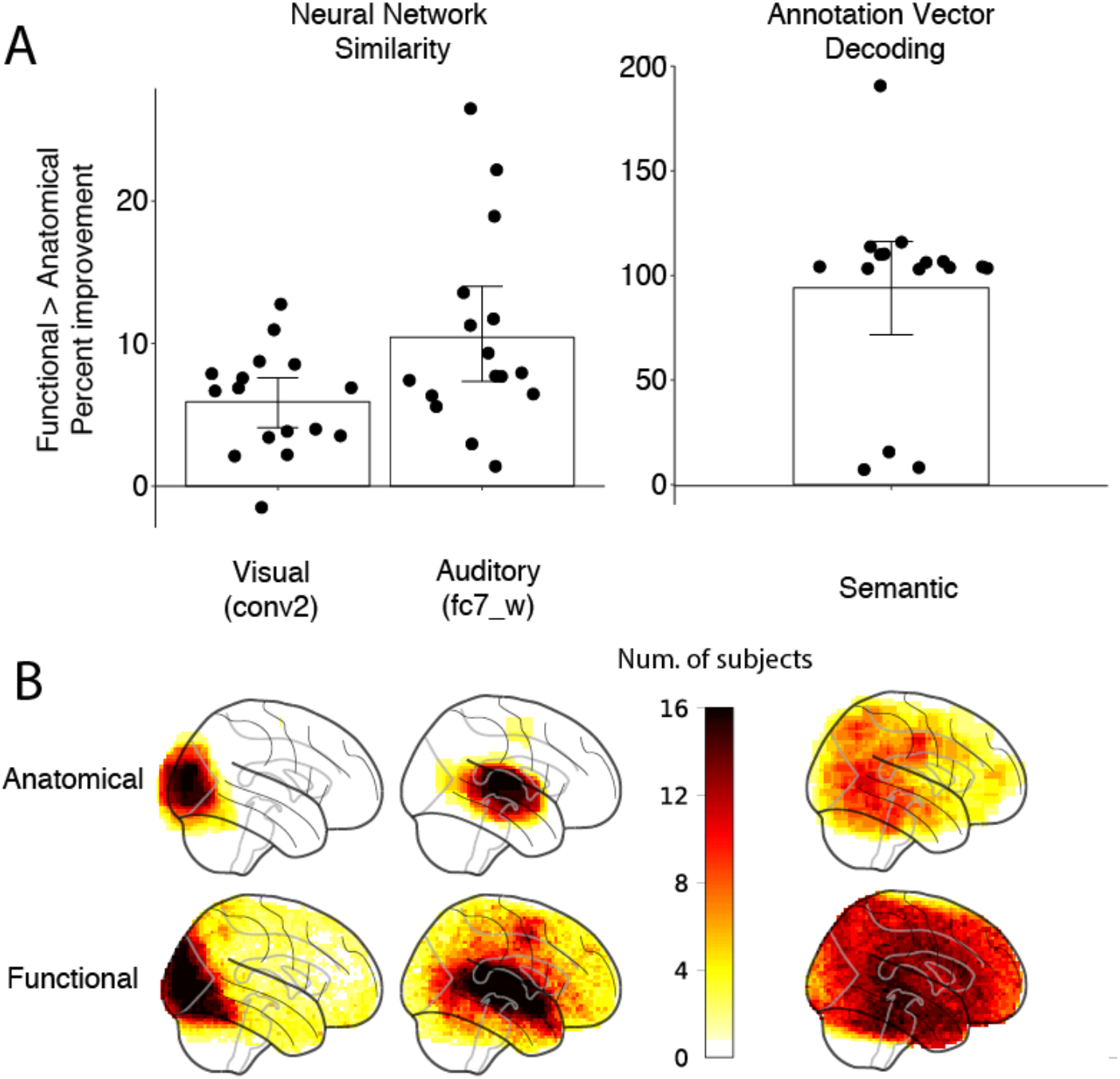
In each subject, we calculated the percent improvement of functional searchlight over anatomical searchlight for the top-performing 1% of searchlights from each approach. (A) Each dot represents a subject’s percent improvement for example layers in the AlexNet visual network (conv2) and KellNet auditory network (fc7_W), as well as for annotation vector decoding. Error bars depict 95% CIs calculated from bootstrapping. Absolute performance levels for each searchlight type and non-parametric baselines can be found in Figure S3. (B) We visualize which voxels contained model-based information by depicting the count of the number of subjects for whom that voxel contributed to one or more of the top 1% of their functional and anatomical searchlights.

To generalize these findings beyond sensory systems, we then analyzed the representation of semantic content in the brain. Theories of semantic cognition[13] and recent findings[14] suggest that such content is widely distributed, and yet the extent may have been underestimated empirically with current methods. We decoded semantic content of each time-point in the movie by predicting sentence embeddings of the movie’s scene annotations from brain activity[15]. We found considerably better decoding of annotation embeddings for the top 1% of functional searchlights than the top 1% anatomical searchlights (Figure 2A right). In other words, aggregating information that is anatomically distributed throughout the brain (Figure 2B right) provided a more accurate representation that was beneficial in probing neural representations of semantic content.

We performed follow-up analyses to explore the parameters that might affect functional searchlight performance and what might account for its consistent advantage over the anatomical searchlight, including: the searchlight radius (Figure S4), the role of bilateral information (Figure S5), and the number of brain voxels per searchlight (Figure S6).

To confirm that running a searchlight in functional space can capture distributed information throughout the brain, we performed the RSA described above on simulated data with a known spatial signal distribution. In particular, we simulated fMRI data[16] where the signal from the units in the first fully connected layer of AlexNet was inserted into random voxels with varying degrees of spatial smoothness. Our results indicated that the relative improvement afforded by the functional searchlight is best when the signal is highly distributed throughout the brain (Figure 3). This supports our interpretation that functional searchlight out-performs anatomical searchlight because it picks up on information relevant for perceptual and cognitive processing that is distributed throughout the brain.

**Figure 3.**
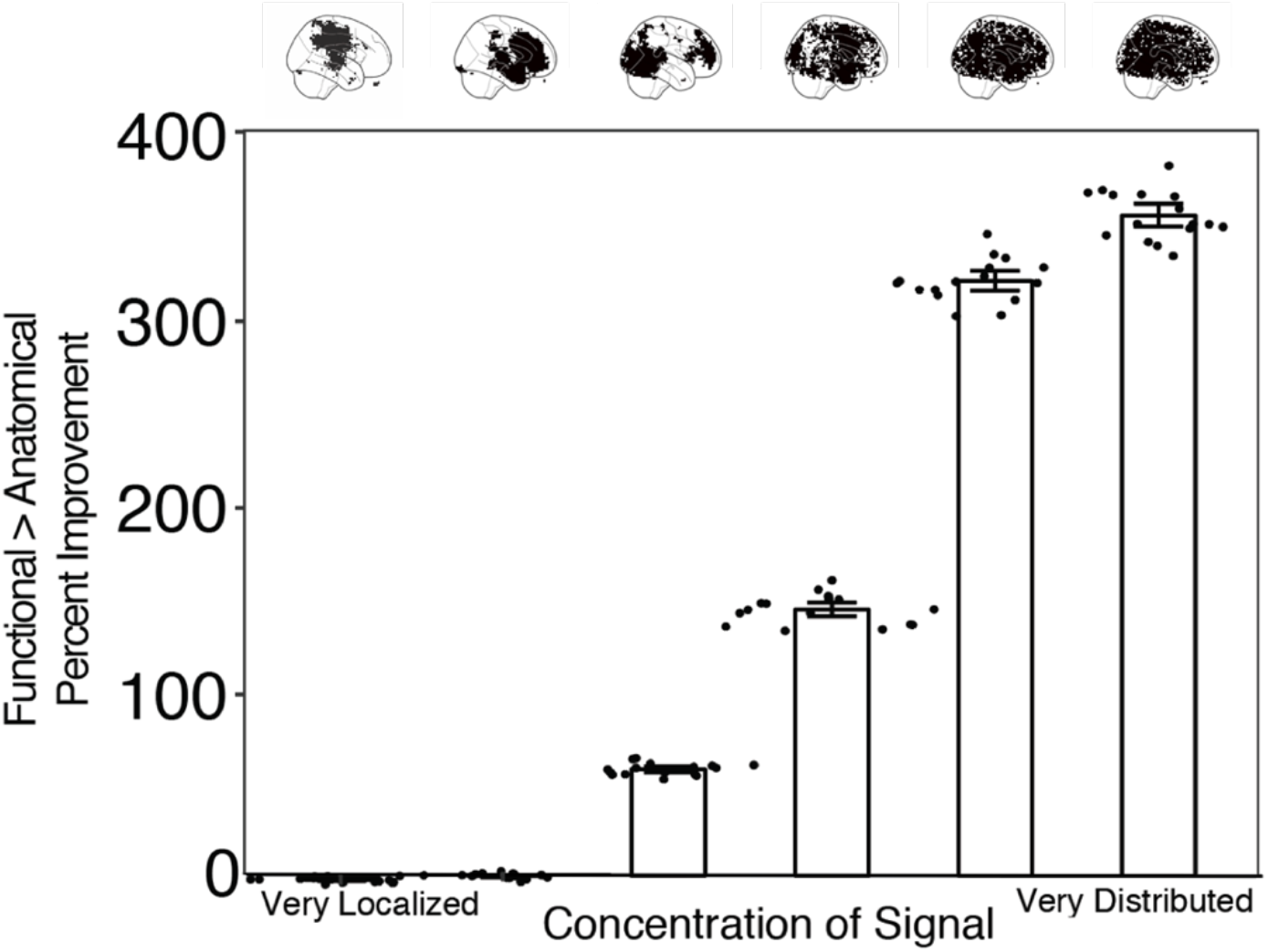
We simulated fMRI data with varying degrees of localized signal. Signal localization was varied by sampling locations of signal voxels from a Gaussian Random Field (GRF) and varying its FWHM (the x-axis in this figure). In particular, a GRF is simulated on the brain and the location of the signal voxels is determined by the highest values of the GRF. A GRF with high FWHM is smoother, so large values will tend to be clustered together. Therefore, for a GRF with high FWHM, the *n* highest values will be localized together. These highest values get more distributed as the FWHM decreases. We ran our RSA analysis for the first fully connected layer of AlexNet (see Methods). The error bars show 95% bootstrapped CIs and the dots represent individual subject improvements. As the signal transitioned from localized to distributed, the relative gain in performance of the functional searchlight increased.

Searchlight analysis, a common multivariate technique in fMRI based on anatomical volumes, is biased toward finding localized information. The functional searchlight is a way to better characterize distributed representations without assumptions about anatomical locality. This approach revealed a tighter correspondence between the human brain and computational models. An important methodological conclusion of this work is that conversion to functional space is worthwhile as a preprocessing step prior to searchlight analysis (if aligning data for SRM are available, like a movie). By enhancing model-based analysis in this way, fMRI can inform the development of new, biologically plausible computational frameworks of cognitive function. This work also has theoretical implications in suggesting that, during viewing of a naturalistic audiovisual stimuli, the brain’s representation of visual, auditory, and semantic information encoded within deep neural network and natural language processing models is more widely distributed than previously thought.

## Methods

### Participants

Full details can be found in the original publication of this dataset[9] and on the data sharing website: https://dataspace.princeton.edu/jspui/handle/88435/dsp01nz8062179. We included the 16 participants who viewed the entire movie. They were right-handed, had normal or corrected-to-normal vision, and had never seen BBC’s Sherlock previously. Informed written consent was obtained in accordance with a protocol approved by the Princeton University IRB.

### Materials

Participants watched a 48-minute clip of the BBC television series “Sherlock” from Season 1, Episode 1. The stimulus was divided into two similarly sized segments (part 1 and part 2). At the beginning of each segment, an audio-visual cartoon was displayed for 30 s. Every time point of the movie was coded manually for various content categories[9]. These include which character was in focus of the camera, which character was speaking, whether the scene was indoor or outdoor, and whether there was music playing.

### Data acquisition and preprocessing

The fMRI data were collected on a 3-T scanner (Siemens Skyra) with a 20-channel head coil. A T2*-weighted echo-planar imaging (EPI) pulse sequence (TE 28 ms, TR 1500 ms, flip angle 64°, 27 slices, 4 mm thickness, 3 × 3 mm in-plane resolution, FOV 192 × 192 mm, whole-brain coverage) was used to acquire functional images. A T1-weighted MPRAGE pulse sequence (0.89 mm isotropic resolution) was used to acquire anatomical images. Preprocessing included slice-time correction, motion correction, linear detrending, high-pass filtering (140 s cutoff), and registration of the functional volumes to a template brain (MNI) at 3mm resolution. Every voxel was z-scored in time within movie run and timing was aligned across participants.

### Searchlight analysis

Searchlight analysis was performed by iterating the same computation over subset volumes of voxels across the brain. The BrainIAK package was utilized for parallelizing the computations [17]. Each searchlight was a tensor centered on every voxel inside the brain, with a radius of 3 voxels. This tensor was thus 7 × 7 × 7 × *t* such that it represented a cube of voxels and *t* time points from the movie (*t*=946 or 1030 for part 1 and part 2, respectively). When near the edge of the brain, anatomical searchlights often contained fewer than the full 343 voxels of the cube because non-brain voxels were excluded (minimum = 115). In the resulting brain map, each voxel’s value represents the output of the analysis on that searchlight’s voxels. In this study, we refer to this traditional searchlight analysis as an “anatomical searchlight”. We use the anatomical searchlight as a baseline to compare with our new functional searchlight approach.

In the functional searchlight, data were first embedded into a space where distances between voxels were based on functional distance rather than anatomical distance, and then a searchlight was run on these transformed data. Voxels that represent similar content will be close to one another in functional space and thus included in the same searchlight. To create this functional embedding, we use shared response modeling[8] (SRM), which maps voxels into a lower-dimensional feature space of fMRI activity that reflects what is shared across participants while they view a common stimulus. SRM is an unsupervised dimensionality reduction technique used for functionally aligning multiple subjects’ fMRI data together on a dimensionality-reduced shared space. Let *X_i_* denote a *v* (voxels) by *t* (timepoints) matrix that represent a single subject *i*’s data. *k* is the reduced number of dimensions or features to which these data will be mapped. SRM learns *n* (number of participants) orthogonal matrices *W_i_* of shape *k* by *v* such that the quantity 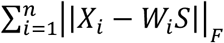 is minimized, where *S* is the *k* by *t* matrix representing the shared response. The resulting *W_i_* for each subject is a weight matrix that can be used to transform the subject’s fMRI data in voxel space into a lower-dimensional shared space.

SRM can be applied to data in which participants are engaged by a common stimulus. In our case, all subjects viewed the same audiovisual stimulus for a total of 1973 TRs. We wanted to train the SRM on a separate portion of the data from what was used for the main analyses. We thus divided the data into two parts, the first with 946 TRs (part 1) and the second with 1030 TRs (part 2). We trained the SRM on part 1 and ran our analyses on part 2, then reversed this order to train the SRM on part 2 and run analyses on part 1. We averaged the results of both analysis folds.

SRM is used to create a *k*-dimensional shared feature space, where *k* is selected via time-segment matching (see below). This means that each voxel had a weight, from their subject-specific orthogonal weight matrix, for how much that voxel loads on to each of the *k* features. These weights were then used as coordinates to remap that voxel into a *k*-dimensional space, independent of the voxel’s anatomical location. Voxels with a weight of 0 for each of the *k* features were discarded, due to them not contributing at all to the shared feature space. We then defined a searchlight for each voxel in functional space as the *j*-nearest neighbors of that voxel, where distance was defined by the cosine distance between voxels’ functional space coordinates. We chose *j* to match the size of the anatomical searchlights (in this case, *j*=342).

### Time-segment matching

We ran a time-segment matching analysis[8] to select the number of dimensions for the functional space. An SRM was fitted on one half of the training data (i.e., one quarter of the episode) and used on the second half of the training data in a leave-one-participant-out cross-validation. Time segments of 14s were taken from the left-out participant’s data and correlated with all segments of the same length in the shared space averaged over the remaining participants. The time-segment matching accuracy for each subject was the proportion of test time segments that were maximally correlated with the correct time segment in the shared space. We varied the number of features from 3 to 400 (step size of 10) and observed that accuracy plateaued in both parts around 200 features (Figure S1). Given these results, we used *k*=200 dimensions for the functional space.

### DNN similarity analysis

We compared the similarity of activity patterns within functional and anatomical searchlights to the similarity of visual and auditory representations derived from deep neural network models. In particular, we compared a time-point by time-point RSA of searchlight data to the similarity structure of visual (AlexNet[11]) and auditory (KellNet[12]) DNNs also viewing the movie. AlexNet is a visual object recognition network with five convolutional layers (conv1-5) and three fully connected layers (fc6-8). It takes a 227×227×3 colored image and outputs a 1000-unit vector in its last fully connected layer, which contains its confidence of the image belonging to one of 1000 image categories. KellNet is an auditory network with separate branches for word recognition and music genre recognition. It takes as input a cochleagram generated from an audio waveform and returns a word category and music genre category in each respective branch. The two branches share three convolutional layers (conv1-3) and then each have their own two convolutional layers (conv4/5_W for word recognition and conv4/5_G for music genre recognition) as well as their own two fully connected layers (fc1/top_W for word recognition and fc1/top_G for music genre recognition). fctop_W is a 587-unit vector that contains confidence in the sound belonging to one of 587 word categories. fctop_G is a 41-unit vector that contains confidence in the sound belonging to one of 41 music genres. In order to keep the naming convention consistent with AlexNet, we refer to fc1_W and fc1_G as fc6_W and fc6_G, respectively, along with fctop_W and fctop_G as fc7_W and fc7_G, respectively.

To obtain the model-based activity for the visual modality, AlexNet received individual frames of the movie. We extracted the activations from each convolutional layer and fully connected layer. We averaged the activity across a subset of frames within the same TR so that the output was a pattern of activity for each TR of the movie. Each TR lasted 1.5 seconds and the movie had 25 frames per second, so a single TR theoretically contained 37.5 frames. We averaged activity using the middle 25 frames of each TR to reduce the autocorrelation across DNN features of different TRs. The end result was AlexNet activity of the entire Sherlock movie in TR intervals. To obtain the model-based activity for the auditory modality, we broke up the movie into 1.5s audio clips (to match the TR) and generated cochleagrams for each TR to be used as an input to KellNet. The activity these clips generated at each of the convolutional layers was then stored. This model-based activity was used to create a representational similarity matrix (RSM) for each layer. Let *t* be the number of TRs in the movie. For each layer and for each model, we constructed a *t* × *t* correlation matrix by correlating unit activity vectors across different TRs.

For each subject, we also constructed an RSM within each searchlight of the brain by correlating voxel activity across different TRs. In these brain RSMs, elements close to the diagonal will be highly similar due to the autocorrelation of the BOLD response. To remove this potential confound in measuring representational similarity, we imposed a buffer of 10 TRs (15s), such that we only retained correlations between TRs separated by at least this much time. These retained off-diagonal elements (i.e., TR pairs) of the model RSM and brain RSM were unraveled into vectors of the same length. We then correlated these vectors to quantify the similarity between representations in each searchlight and in a given layer of a computational model, and assigned this second-order correlation to the center voxel.

### Natural language decoding analysis

We compared functional and anatomical searchlight performance on a natural language decoding analysis[15]. The Sherlock dataset contains a handwritten annotation of what is happening in each TR of the movie (one sentence per TR, averaging 18 words). In previous work, Natural language processing (NLP) techniques were used to create semantic vector representations of each annotation[15]. That is, each TR was represented by a 100-dimensional vector that encodes the semantic meaning of the corresponding annotation. These vectors were constructed from word embeddings built using a latent-variable modeling approach[15,18] on the Wikipedia corpus for each word in a given annotation. A domain-specific re-centering was applied to these word embeddings to make each of them discriminative within the average topic of the Sherlock annotation vocabulary. An embedding was defined for the entire annotation as a weighted average of the constituent word embeddings, with weights determined by relative word frequencies.

We predicted these annotation vectors from brain activity using a “scene ranking” analysis[15]. We used the first half of the movie to train the SRM and create the functional space, which was then applied to the second half of the movie prior to analysis. Within the second half, we trained a ridge regression model on half of the timepoints (i.e., quarter of the movie) to take searchlight voxels and predict the 100-dimensional annotation embedding of the other half of timepoints. Specifically, we trained the SRM and learned the functional space on part 1, then trained the ridge regression model on the first 530 TRs of part 2 to predict the last 500 TRs of part 2. This resulted in a predicted 100-dimensional annotation vector for each of these 500 TRs, which could be compared against the actual annotation vectors for these TRs to quantify performance. We divided the 500 predicted and actual annotation vectors into 25 evenly sized bins, comprising 20 annotation vectors from consecutive TRs. We then computed a 25 × 25 correlation matrix *M*, where *M_ij_* denotes the correlation of predicted bin *i* with the actual bin *j*. Our reported accuracy is the proportion of predicted bins that were most highly correlated with the corresponding actual bin (chance accuracy = 1/25 or 4%). Unlike previous analyses, we did not reverse the order and train SRM on the second part and do analysis on the first part. This is because the annotation vectors we used from [15,18] were normalized using data from part 1 of the movie, so to avoid double dipping, we refrained from doing annotation vector decoding on part 1 of the movie.

### Statistics

For our primary analysis (Figure 2), we obtained the average performance of the top 1% of searchlights for both the functional searchlight approach and the anatomical searchlight approach. We then subtracted the empirically derived chance performance for each searchlight type (see below) from these averages and calculated the percent improvement of functional searchlight over anatomical searchlight. To test significance, we generated 95% confidence intervals of percent improvement by bootstrap resampling across participants[19]. Specifically, we sampled with replacement from the percent improvements of our 16 subjects, calculated the average of each sample, and repeated the process 10,000 times. The logic of this approach is that to the extent that the effect is reliable across participants, the participants should be interchangeable and it should not matter which subjects are sampled on a given iteration. We report the 95% CIs as the 2.5^th^ percentile and 97.5^th^ percentile of the resampled distribution. Functional searchlight performed significantly better than anatomical searchlight if and only if the lower bound of the confidence interval for percent improvement is above 0.

We empirically determined the chance level using a “rolling analysis.” Across 100 iterations, we misaligned the brain and model data in time by shifting one of them with respect to the other in multiples of 10 timepoints. By repeating all analyses after such misalignment, we populated a null distribution of performance. We defined chance performance for a given subject and analysis as the mean of this null distribution. This was then used as a reference for the real value, to calculate performance above chance.

### Simulation

We assume that functional searchlight outperforms anatomical searchlight because it can capture distributed information. We evaluated this interpretation by simulating fMRI data that varied in how broadly information was distributed across the brain and repeating the AlexNet analysis. The simulation was performed using the fmrisim package in BrainIAK[16]. In particular, we simulated realistic fMRI noise from participant templates [20] and chose a set of voxels in the simulated brain to insert signal. We simulated 16 participants whole-brain data and inserted signal from the 4096 units of fc6 in AlexNet into 4096 voxels. Hence, if there were voxels in the brain that responded exactly the same the fc6 layer of AlexNet then their activity would look something like the signal that was simulated. This signal was generated by convolving the unit activity (averaged across frames) for each TR with a double-gamma hemodynamic response function. This signal was then added to the realistic neural noise so that the overall timeseries contained signal from the deep network, while retaining properties of BOLD such as autocorrelation and lag. The magnitude of the inserted signal was set at 0.5 percent signal change.

The voxels chosen to carry signal were determined based on how distributed the induced signal was. In particular, we simulated a Gaussian Random Field (GRF) the same shape as each simulated participant’s brain and chose the voxels containing the highest 4,096 GRF values as the locations of the signal voxels. The GRF has a specific smoothness, parameterized by the full-width half max (FWHM). If the GRF is smooth, then all of the highest values are likely to be localized in a single area. However, if the GRF is not smooth, then these signal voxels will be distributed throughout the brain. We created volumes with signals distributed over FWHMs of 0.5, 1.0, 2.0, 4.0, 8.0, 16.0. In other words, we simulated participant data with signal distributed locally or broadly throughout the brain. To quantify performance, we repeated the main representational similarity analysis of AlexNet on simulated data from anatomical and functional searchlights.

### Code availability

Custom computer code was written to implement the analyses described in the manuscript. This code is available at this link: https://github.com/sreejank/DistributedCodingBrain

## Acknowledgments

The authors thank Janice Chen and her colleagues in the laboratories of Ken Norman and Uri Hasson at Princeton University for collecting and sharing the dataset analyzed herein. We also thank Ken Norman for helping us obtain the annotation embeddings. This work was supported by NSF grant CCF 1839308 and BCS 1558497, NIH grant R01 MH069456 and R01 MH108591, and the Canadian Institute for Advanced Research.

## Figure Supplements

**Figure S1.**
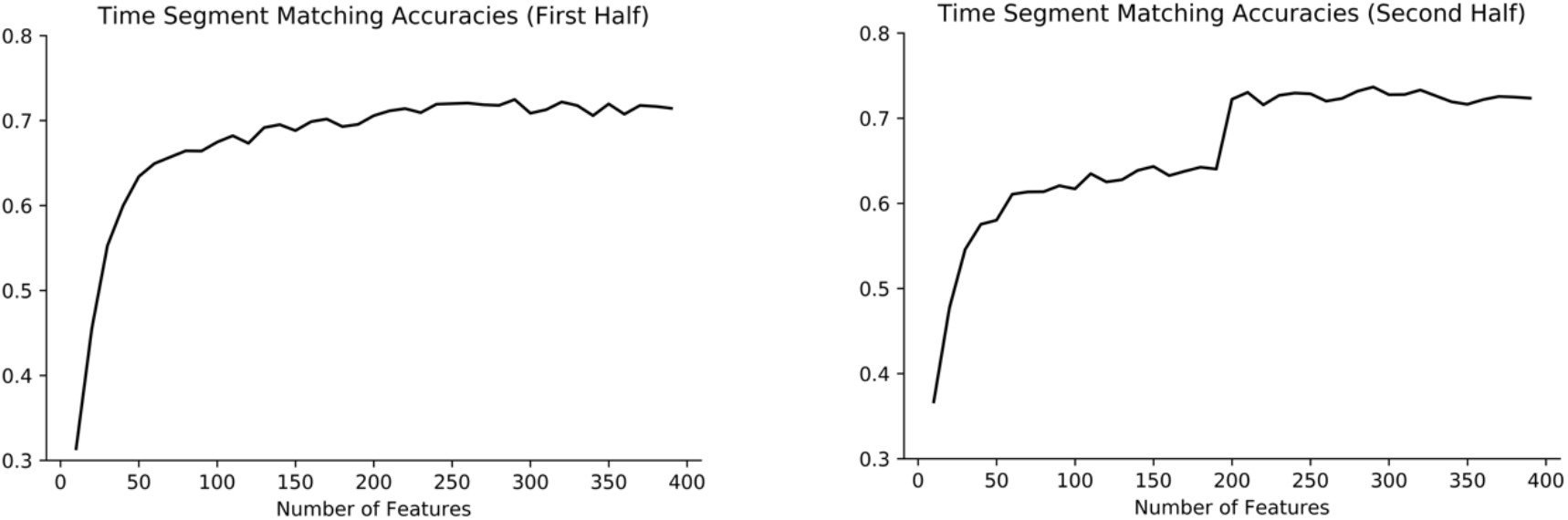
We performed a time-segment matching analysis[8] to determine how many SRM dimensions to use. Within the training half of the movie, we trained an SRM with the corresponding number of dimensions (*x* axis) on one half of the TRs and tested on the second half of the TRs. The goal was to predict from which time window in the movie the test data were obtained (chance = 0.0212). Time-segment matching proportion correct (*y* axis) plateaued at approximately 200 dimensions considering both folds of the overall analysis.

**Figure S2.**
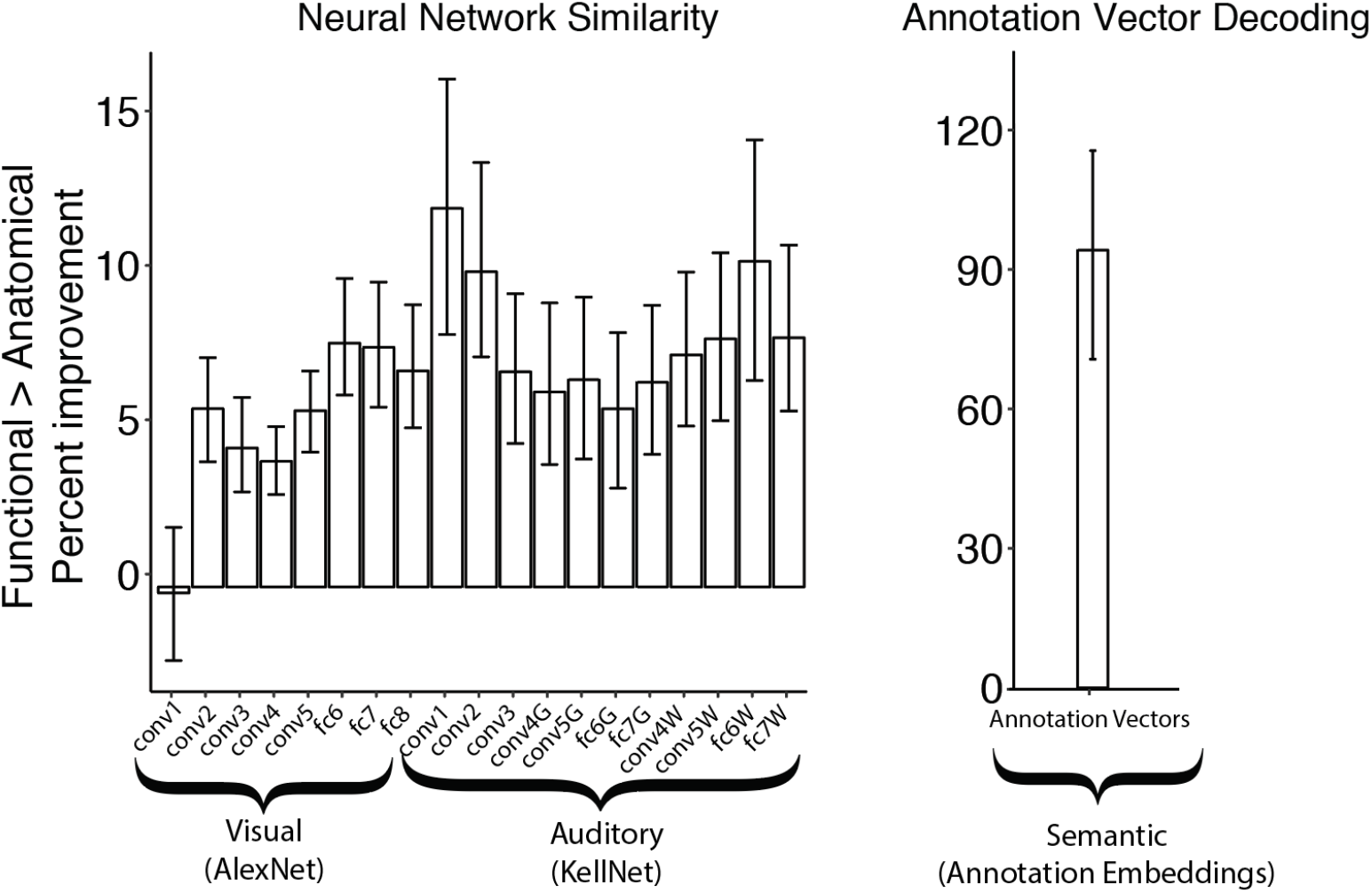
Functional searchlight percent improvements (error bars denote 95% CIs) within the top 1% of searchlights for both DNN RSA analyses (for all layers) and annotation vector decoding analyses. Functional searchlight performed significantly better if the lower 95% confidence bound was above 0 (all cases except AlexNet conv1).

**Figure S3.**
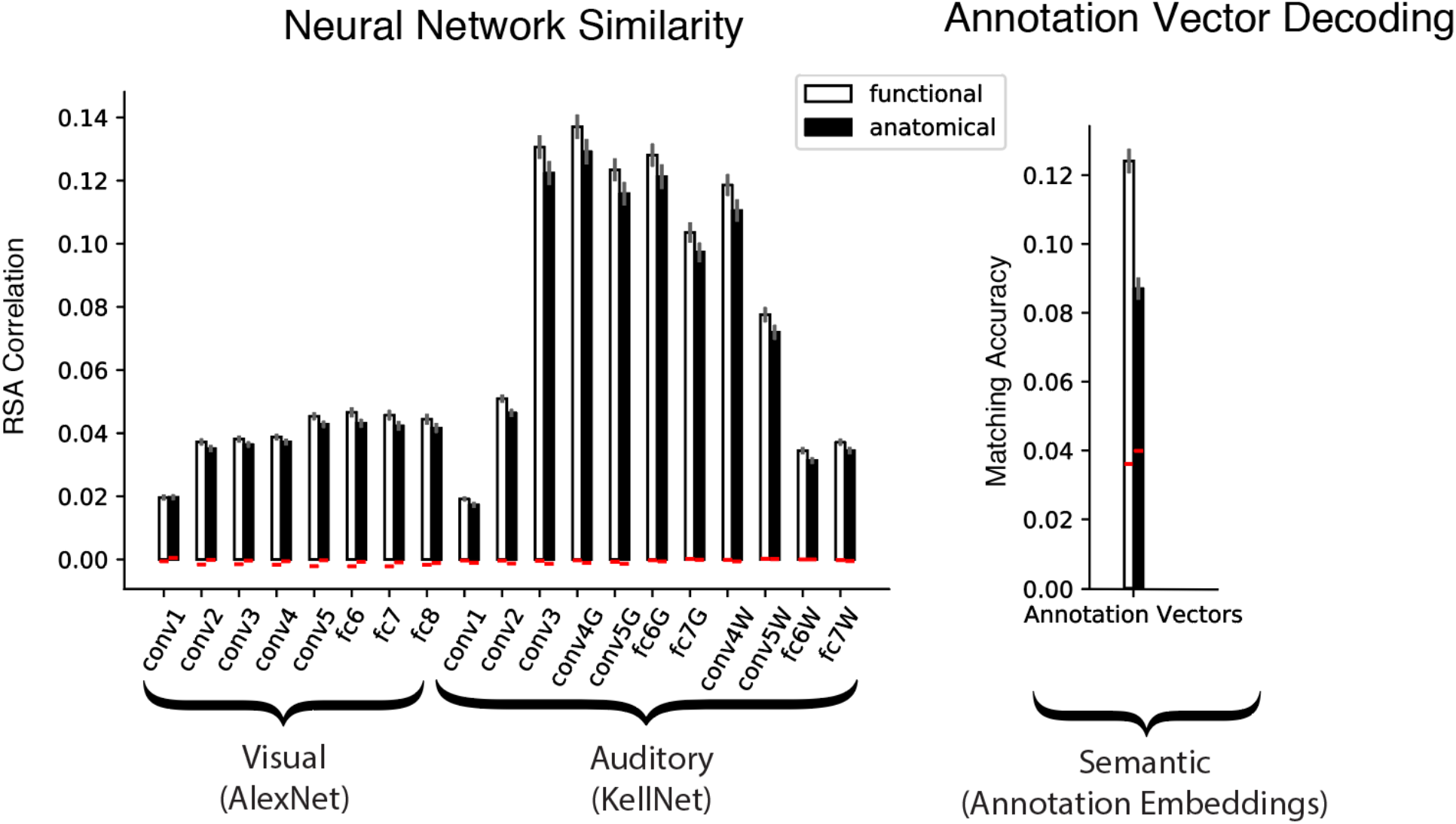
Average performance in the top 1% of functional and anatomical searchlights for neural network similarity and semantic vector decoding. Error bars represent standard error across subjects. To calculate percent improvement as used in the main analyses (Figures 2 and S2), we subtracted chance performance (mean of null distribution estimated non-parametrically by rolling data in time, shown here as the red bars) from the performance of each searchlight type and calculated the percent improvement from anatomical to functional. Note that the neural network data (left) are RSA correlations while the annotation embedding data (right) are decoding accuracies.

**Figure S4.**
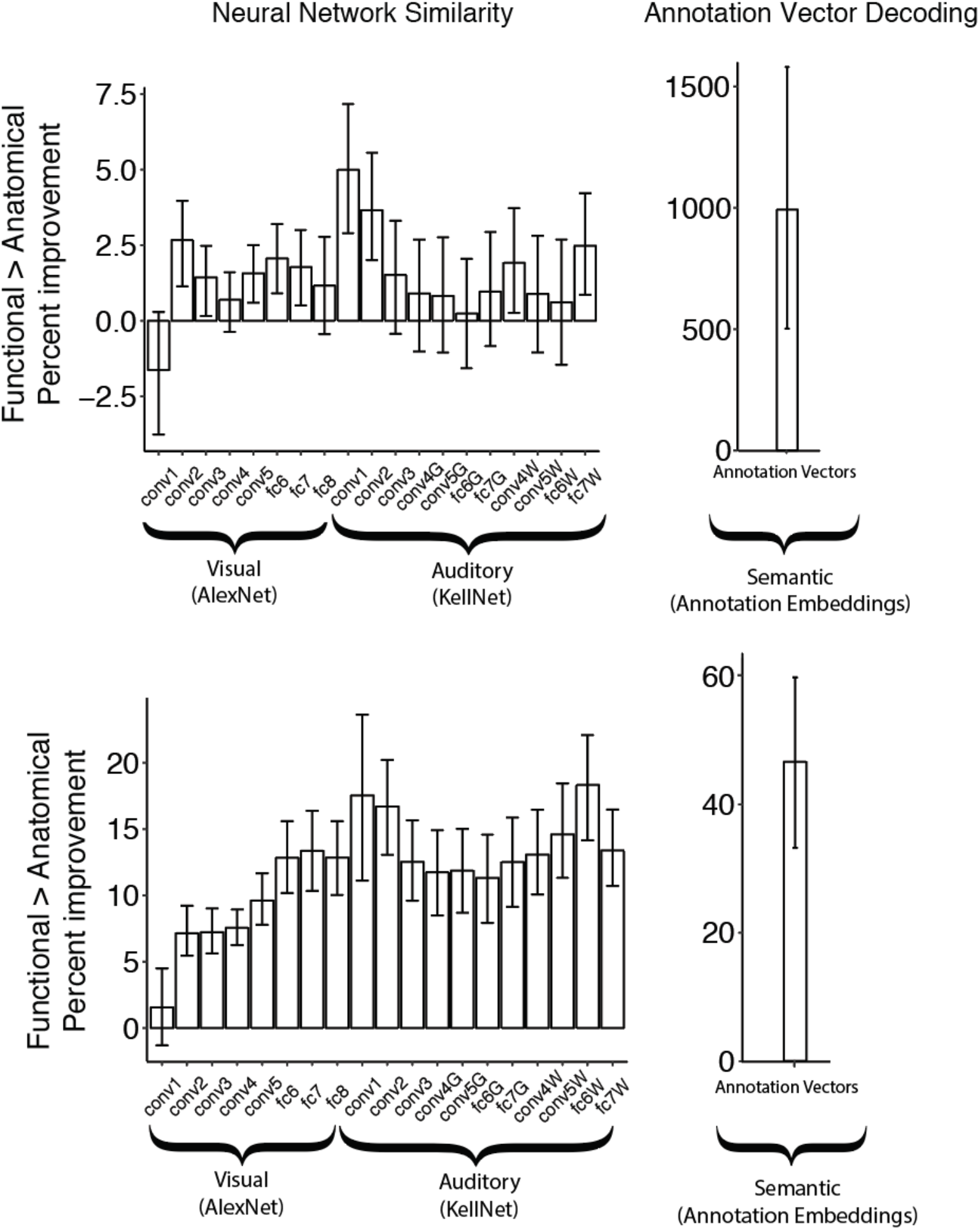
For our main analyses, we chose a searchlight radius = 3 (number of voxels for each searchlight 7 × 7 × 7 = 343). Here we report results for radius = 2 (top row; number of voxels: 5 × 5 × 5 = 125) and radius = 4 (bottom row; number of voxels: 9 × 9 × 9 = 729). A larger radius led to a greater advantage for functional over anatomical searchlights in neural network analyses and a smaller advantage for the NLP analyses.

**Figure S5.**
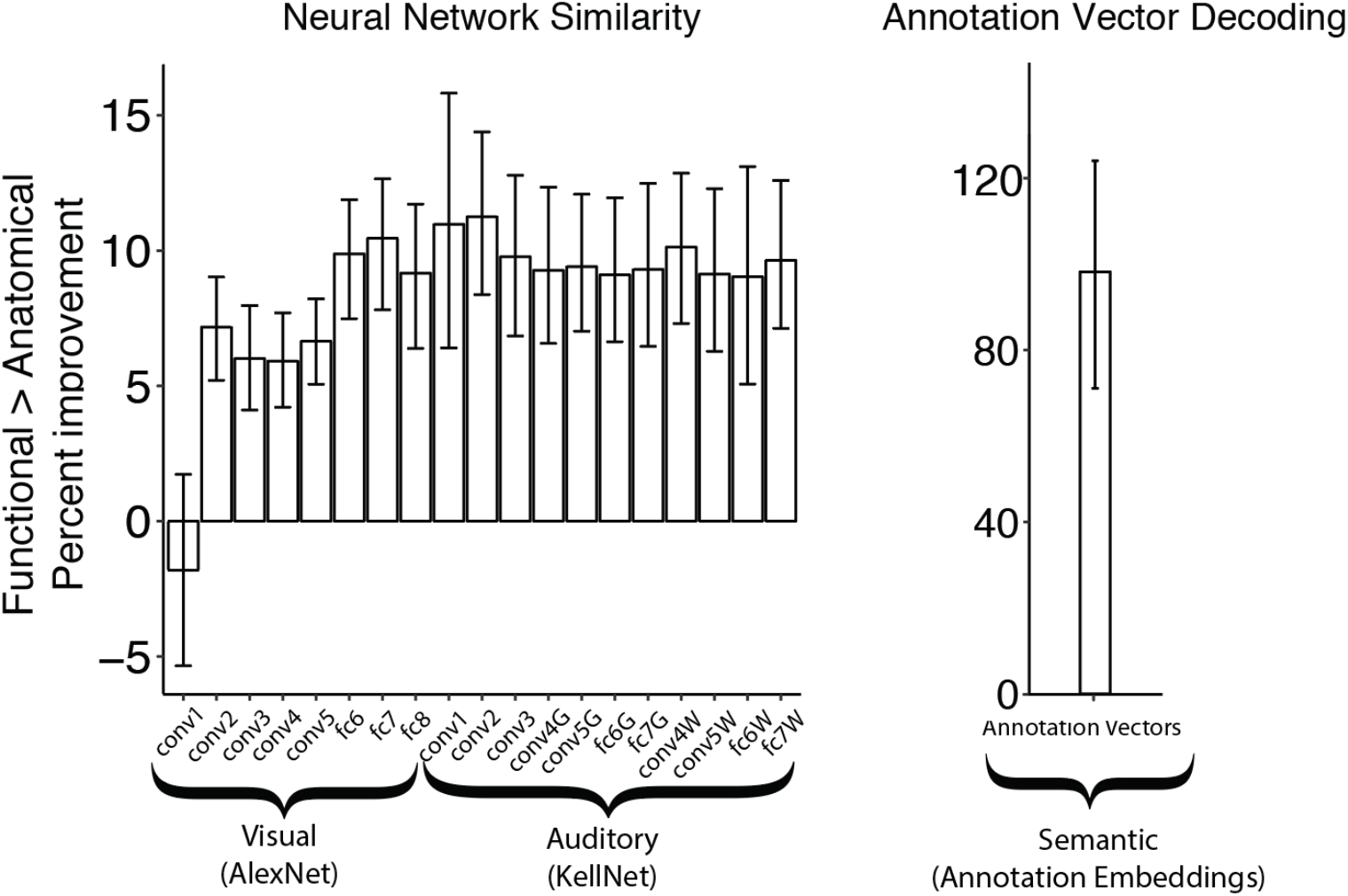
To test whether the improved performance of functional searchlights reflects their ability to aggregate information across hemispheres from bilateral regions that are anatomically distant but functionally homologous (e.g., left and right auditory cortex), we re-ran all of our analyses within one hemisphere of the brain at a time (including fitting SRM) by completely ignoring the other hemisphere. The error bars show 95% bootstrapped CIs. Even in the absence of bilateral information, functional searchlight still consistently outperforms anatomical searchlight.

**Figure S6.**
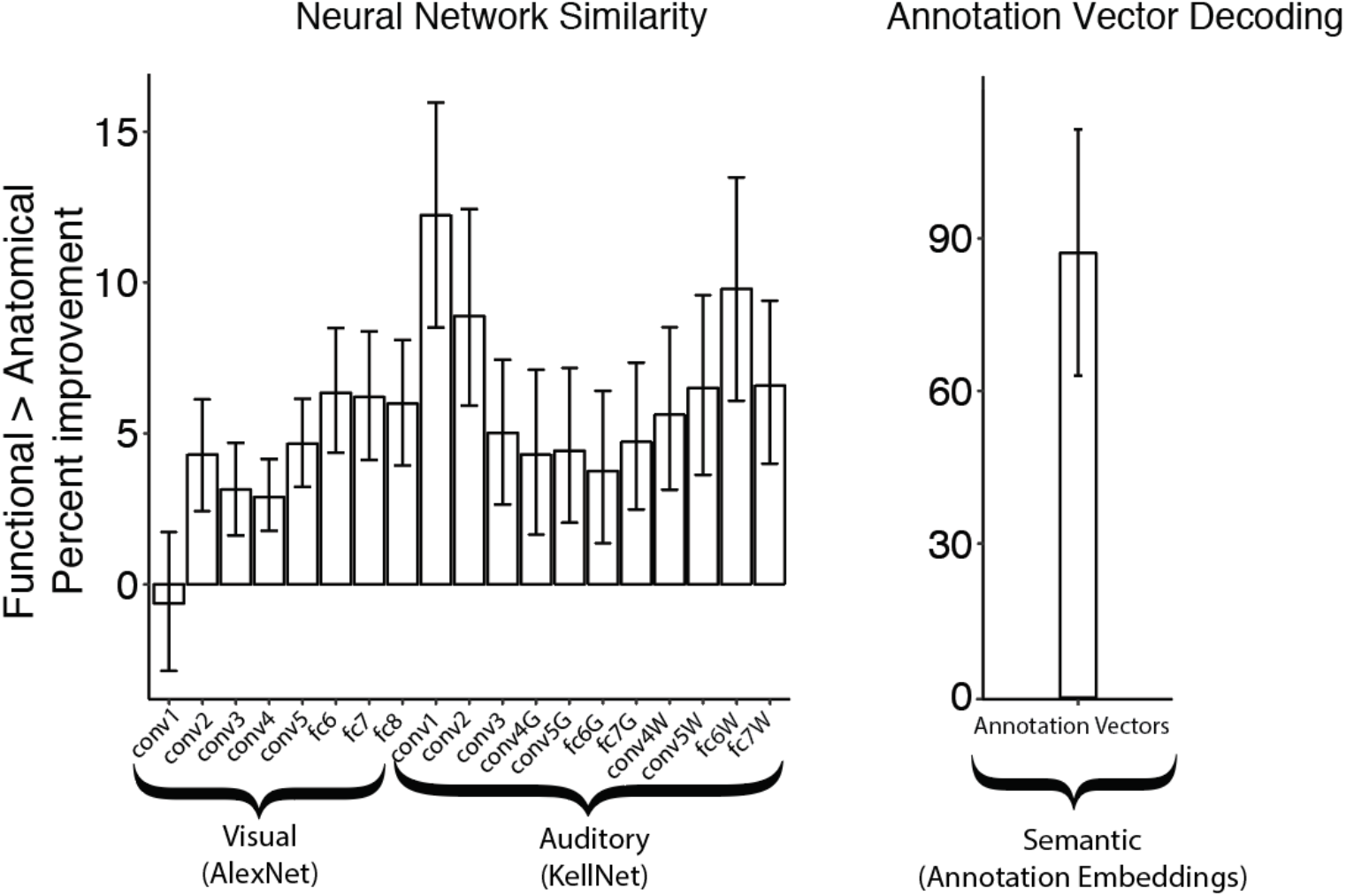
Some anatomical searchlight volumes on the edge of the brain would contain non-brain voxels that need to be excluded, resulting in fewer brain voxels defining the activity pattern. Since we are taking the nearest neighbors in functional space, the number of voxels per functional searchlight is always the same. To ensure functional searchlight is not benefitting from having more voxels in certain searchlights, we ran a version of our analyses where each functional searchlight was forced to contain the exact same number of voxels as the anatomical searchlight. The error bars show 95% bootstrapped CIs. Even when we force the number of voxels per searchlight to match, functional searchlight still performs better.

